# Disruption of the KH1 domain of *Fmr1* leads to transcriptional alterations and attentional deficits in rats

**DOI:** 10.1101/338988

**Authors:** Carla E. M. Golden, Michael S. Breen, Lacin Koro, Sankalp Sonar, Kristi Niblo, Andrew Browne, Daniele Di Marino, Silvia De Rubeis, Mark G. Baxter, Joseph D. Buxbaum, Hala Harony-Nicolas

## Abstract

Fragile X Syndrome (FXS) is a neurodevelopmental disorder caused by mutations in the *FMR1* gene. FXS is a leading monogenic cause of autism spectrum disorder (ASD) and inherited intellectual disability (ID). In most cases, the mutation is an expansion of a microsatellite (CGG triplet), which leads to suppressed expression of the fragile X mental retardation protein (FMRP), an RNA-binding protein involved in multiple aspects of mRNA metabolism. Interestingly, we found that the previously published *Fmr1* knockout rat model of FXS expresses a transcript with an in-frame deletion of a K-homology (KH) domain, KH1. KH domains are RNA-binding domains of *FMR1* and several of the few, known point mutations associated with FXS are found within them. We observed that this deletion leads to medial prefrontal cortex (mPFC)-dependent attention deficits, similar to those observed in FXS, and to alterations in transcriptional profiles within the mPFC, which mapped to two weighted gene coexpression network analysis modules. We demonstrated that these modules are conserved in human frontal cortex, are enriched for known FMRP targets and for genes involved in neuronal and synaptic processes, and that one is enriched for genes that are implicated in ASD, ID, and schizophrenia. Hub genes in these conserved modules represent potential targets for FXS. These findings provide support for a prefrontal deficit in FXS, indicate that attentional testing might be a reliable cross-species tool for investigating the pathophysiology of FXS and a potential readout for pharmacotherapy testing, and identify dysregulated gene expression modules in a relevant brain region.

**Significance Statement:** The significance of the current study lies in two key domains. First, this study demonstrates that deletion of the Fmrp-KH1 domain is sufficient to cause major mPFC-dependent attention deficits in both males and females, like those observed in both individuals with FXS and in knockout mouse models for FXS. Second, the study shows that deletion of the KH1 domain leads to alterations in the transcriptional profiles within the medial prefrontal cortex (mPFC), which are of potential translational value for subjects with FXS. These findings indicate that attentional testing might be a reliable cross-species tool for investigating the pathophysiology of FXS and a potential readout for pharmacotherapy testing and also highlight hub genes for follow up.

## Introduction

Fragile X Syndrome (FXS) is a neurodevelopmental disorder that is a leading monogenic cause of autism spectrum disorder (ASD) and the most frequent known form of inherited intellectual disability (ID) [1]. FXS is caused by the loss of Fragile X Mental Retardation Protein (FMRP), which is encoded by the *FMR1* gene on chromosome X [1]. FMRP is involved in the regulation of messenger RNA (mRNA) translation [2], shuttling of mRNA to dendritic spines [3], and stability of mRNA [3]. FMRP interacts with RNA through K-homology (KH) domains and the arginine-glycine box (RGG) [4]. In most FXS cases, Fmrp loss occurs when the unstable trinucleotide CGG repeat at the 5’ untranslated region of *FMR1* expands to above 200 copies, resulting in the hypermethylation and transcriptional silencing of *FMR1* [5]. Notably, both point mutations [6] and deletions [7, 8] within the *FMR1* gene coding sequence have also been reported in a small number of individuals, including missense mutations in the KH domains [9-22] (**Figure 1**).

**Figure 1.**
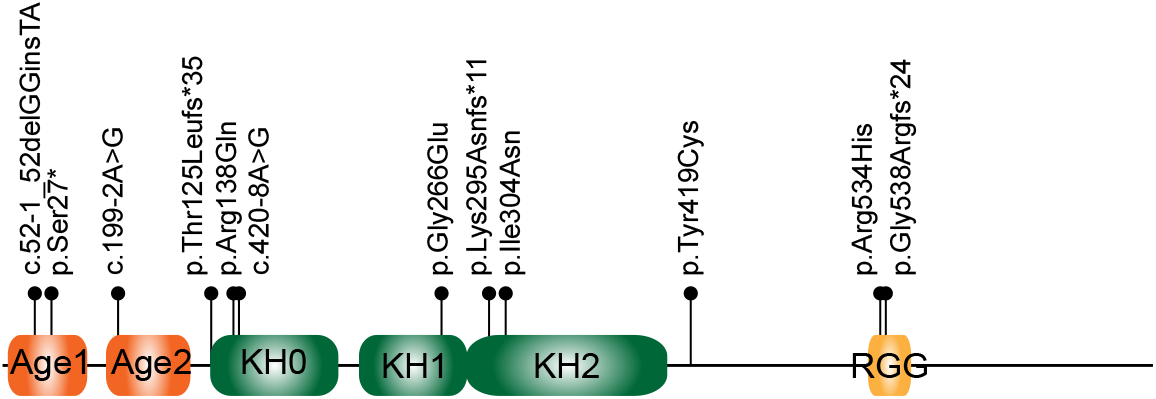
Pathogenic point mutations associated with FXS. All pathogenic mutations in the coding region of the FMR1 gene published [9–22] or deposited in ClinVar are summarized here. The FMRP domains are reported as described in Myrick et al., 2015 [107]. Mutations are indicated using the HGSV nomenclature. Reference sequences used are Q06787 for the protein and NM_002024.5 for the cDNA. The splice-site mutations are indicated by their splice-site nomenclature and localized to the position of the first amino acid predicted to be affected. For c.990+1G>A, we have indicated the amino acid change (p.Lys295Asnfs*11) identified experimentally Quartier et al., 2017 [20]. Age, Agenet-like domain (also known as Tudor domain); KH, K-homology domain; RGG; arginine-glycine-glycine box.

Attention deficits and hyperactivity (ADHD) are very common behavioral manifestations in FXS and are prevalent in both males and females [23]. Attention in rodents requires an intact medial prefrontal cortex (mPFC) [24], which shares functions with prefrontal regions that are anatomically [25–27] and functionally [28, 29] impaired in individuals with FXS. Despite the repeated implication of the mPFC as a nexus of cognitive dysfunction in FXS, it has been the focus of very few pre-clinical studies in animal models [30–32]. In the current study, we used a rat model of FXS to study the involvement of the mPFC in FXS.

To analyze the role of Fmrp in the mPFC, we employed behavioral, molecular, and bioinformatic approaches using a recently described rat model with a 122bp in-frame deletion in exon 8 that was previously shown to have enhanced protein synthesis, exaggerated Group 1 mGluR-dependent long-term depression (LTD), increased spine head width and spine density in the CA1 region of the hippocampus [33, 34], and macroorchidism [33]. We found that this 122bp deletion leads to skipping of exon 8 and the expression of a gene product with loss of the KH1 domain (*Fmr1-^Δ^KH1*). To study how a lack of the KH1 domain affects visuospatial attention, which requires an intact mPFC, we used the five-choice serial reaction time task (5-CSRTT) [35, 36]. In addition, we characterized the transcriptional profiles in the mPFC and identified discrete groups of co-regulated genes.

## Results

### Validation of an *Fmr1-^Δ^KH1* rat model of FXS

Zinc-finger nucleases (ZFN) targeting *Fmr1* were used to introduce a 122bp genomic deletion in the gene in order to develop a rat model of FXS, which has been published on previously [33, 37–40] and referred to here as the *Fmr1-^Δ^KH1* rat (**Figure 2A**). We used both PCR and genomic Sanger sequencing to confirm the 122bp deletion between base-pairs (bp) 18,586 and 18,708 (NCBI reference sequence: NC_005120.4), spanning part of intron 7 and exon 8 of the *Fmr1* gene (**Supplementary Figures 1A, 1B, and 1C**). To examine the effect of the genomic deletion on *Fmr1* mRNA, we amplified the coding sequence between exon 7 and 10 of the *Fmr1* gene using reverse transcriptase polymerase chain reaction (RT-PCR) (**Figure 2B**) followed by Sanger sequencing (**Figure 2C**). This analysis revealed that in the *Fmr1-^Δ^KH1^−/y^* rat at least a portion of mRNAs show a skipping of exon 8, which results in an in-frame deletion (**Figure 2C**). Using Western blot analysis with antibodies directed against the C-terminus or the N-terminus of Fmrp, we detected the full-length Fmrp (~75KDa) in wild type (WT), but not in *Fmr1-^Δ^KH1^−/y^* rats, where we detected a band at a lower molecular weight band (~70KDa) (**Figure 2D**). Immunoprecipitation using a monoclonal against the N-terminus of Fmrp and detection with both a N-terminus and C-terminus directed antibodies (**Supplementary Figure 2A**) recovered the ~75KDa band in WT rats and the 70KDa band in *Fmr1-^Δ^KH1^−/y^* rats. The RNA sequence results, the immunoprecipitation results, and the ~70kDa molecular weight are compatible with an *Fmr1-^Δ^KH1^−/y^* rat in-frame deletion of the 57 amino acids encoded by exon 8 and composing the KH1 domain of Fmrp (**Supplementary Figure 2B**). Notably, the ~70KDa band had decreased expression relative to Fmrp in WT rats, indicating that the in-frame deletion results in reduced synthesis or increased degredation of the resultant protein.

**Figure 2.**
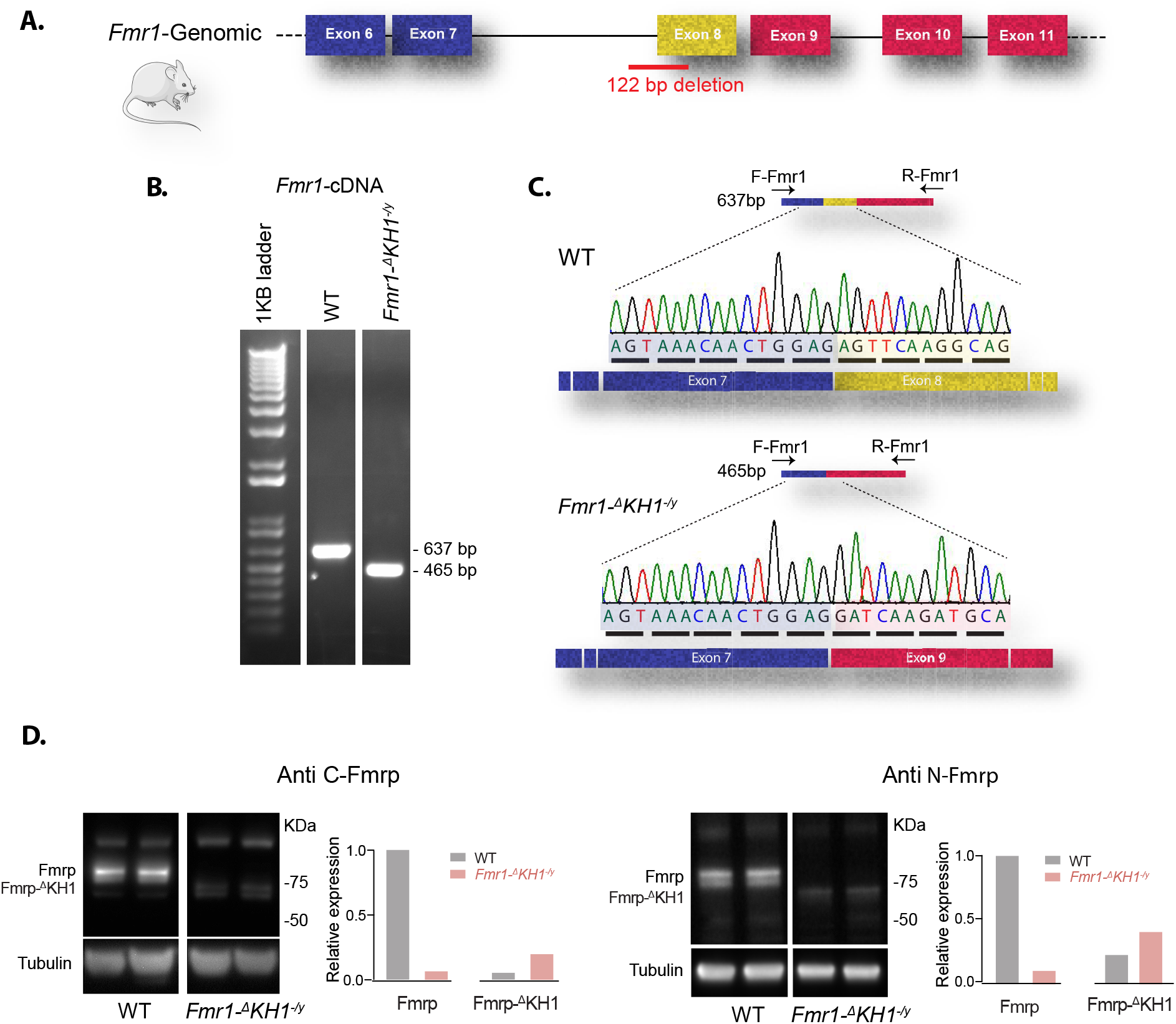
Skipping of exon 8 of the *Fmr1* gene in the *Fmr1-^Δ^KH1* rat model (previously presented as *Fmr1* KO). (**A**) A schematic representation of exons 6-11 of the *Fmr1* gene and the position of the genomic 122bp deletion spanning intron 7 and exon 8 in the *Fmr1-^Δ^KH1* rat model. (**B**) Reverse transcription–polymerase chain reaction (RT–PCR) analysis primed with a forward primer (F-Fmr1), designed to align to exon junction 6/7, and a reverse primer (R-Fmr1), designed to align to exon junction 9/10 (as described in C and detailed in Supplementary Table 3). (**C**) Sanger sequencing of the RT PCR products of WT and *Fmr1-^Δ^KH1^−/y^* rats, primed with the F-Fmr1 primer. Sequencing results confirm the skipping of exon 8 in *Fmr1-^Δ^KH1^−/y^* rats, keeping the frame intact. (**D**) Immunoblotting and quantification of Fmrp and Fmrp-^Δ^KH1 protein levels in WT and *Fmr1-^Δ^KH1^−/y^* mPFC samples, using anti-C-terminus- or N-terminus-Fmrp antibodies.

The FMRP KH1 domain is structurally organized by three anti-parallel β-strands and three α-helices (β1-α1-α2-β2-β’-α’ configuration) with a GxxG loop between α1 and α2 forming a cleft for the RNA binding (**Supplementary Figure 2C**). As shown by the sequence conservation obtained from alignment of *FMR1* orthologs across 58 species [41], the KH1 domain has very strong evolutionary conservation (**Supplementary Figure 2A**). Interestingly, amongst the point mutations in *FMR1* associated with FXS, there are several in the KH domains, including one that lies in the KH1 domain [14] (**Figure 1**). The phenotype associated with this mutation includes the characteristic dysmorphic facial features of FXS, macroorchidism, and ID in comorbidity with ASD and ADHD [14].

The rat with the deletion of KH1 shares some phenotypes with the *Fmr1* knockout (KO) mouse [42–46], including enhanced protein synthesis and alterations in mGluR-dependent LTD and spine density and morphology [33, 34]. We also observed a significant increase in testes:body weight ratio in the *Fmr1-^Δ^KH1^−/y^* rats compared to their WT littermates (**Supplementary Figure 3**; two-tailed T-test, *p* = 0.019), replicating the macroorchidism phenotype reported in the rat [33] and in mouse models for FXS [47]. In contrast to the findings from the *Fmr1* KO mouse model [48–51] however, the rat model does not appear to have increased locomotion in the open field test or impaired spatial memory in the Morris Water Maze task [33, 34]. We replicated this lack of hyperactivity in the open field test, observing a significant effect of time (**Supplementary Figure 4**; repeated measures ANOVA for time, *p* < 0.0001), where animals slowed down over the course of the trial, but no interaction between time and genotype. Using the Barnes maze task, we confirmed the observation that male *Fmr1-^Δ^KH1^−/y^* rats do not have deficits in spatial memory, a hippocampal-dependent form of cognition [34] (**Supplementary Figure 5**). Therefore, we focused our studies on attention, another key cognitive behavior that is impaired in FXS and that is dependent upon a brain region clinically implicated in FXS, the mPFC.

### *Fmr1-^Δ^KH1* rats have deficits in sustained attention

We assessed visuospatial attention with the 5-choice serial reaction time task (5-CSRTT). In this task, rats must respond quickly with a nose poke to briefly presented light stimuli after a five-second delay (**Figure 3A**) [52]. ADHD is comorbid with FXS in both males and females [23]. To compare across sexes, we included hemizygous males (*Fmr1-^Δ^KH1^−/y^*) and homozygous females (*Fmr1-^Δ^KH1^−/−^*), which have comparable genetic vulnerability, as well as heterozygous females (*Fmr1-^Δ^KH^+/−^*). This type of study cannot be carried out in humans due to the fact that homozygous mutations in females with FXS do not exist. Rats were trained in stages where the duration of the light stimulus was incrementally decreased from 60 to 30 to 20 to 10 to 5 to 2.5 s and were progressed to the next stage once performance criterion were met (≥80% accuracy and ≤20% omissions). Decreasing stimulus durations increases demands on sustained attention because briefer stimuli require more attentional effort in order to continue to detect and respond to them successfully.

**Figure 3.**
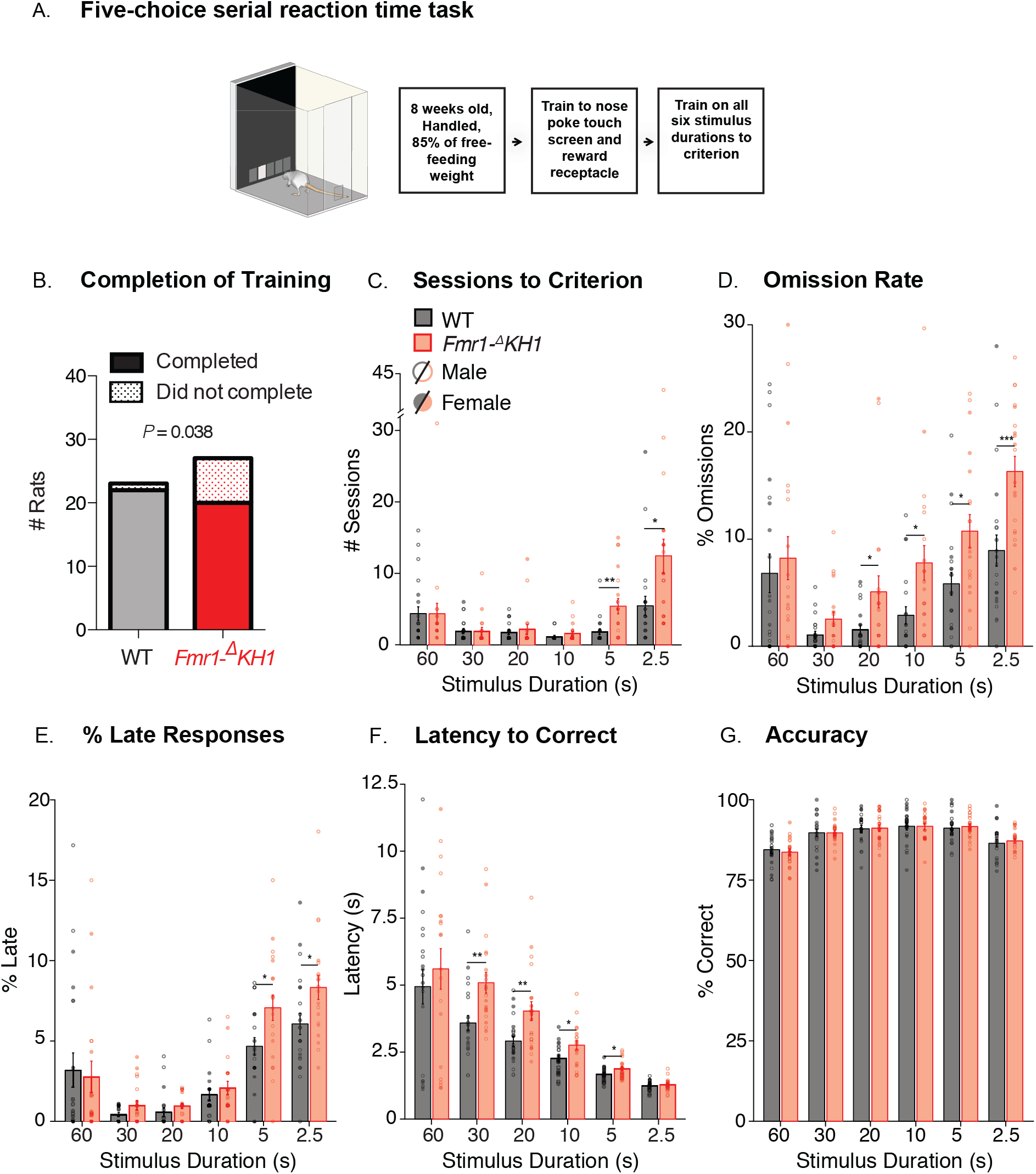
Performance of male and female *Fmr1-^Δ^KH1* rats and WT littermates on the five-choice serial reaction time task (5-CSRTT). (**A**) Schematic of the 5-CSRTT apparatus and training timeline. (**B**) The number of rats that completed training to criterion at a 2.5 s stimulus duration and the reported Phi value from a two by two contingency table that shows a significant association between genotype and completion of training. Across six 5-CSRTT training stages, bars indicate (**C**) mean number of sessions required to reach criterion ± SEM (WT, n = 22; *Fmr1-^Δ^KH1*, n = 20), (**D**) mean percentage of trials that were omitted, (**E**) mean percentage of trials with a late response, (**F**) mean latency to perform a correct response, and (**G**) mean accuracy (# correct / # total responses), black = WT, red = *Fmr1-^Δ^KH1*, open circles = males, filled circles = females, \*\*\**p* < 0.001, \*\**p* < 0.01, \**p* < 0.05.

Two separate analyses were performed: 1) male and female WT rats were compared to male *Fmr1-^Δ^KH1^+/y^* and female *Fmr1-^Δ^KH1^−/−^* rats and 2) female rats were compared amongst each other (WT, *Fmr1-^Δ^KH1^+/−^*, and *Fmr1-^Δ^KH1^−/−^*) (see Methods). Whereas all but one of the WT controls were able to meet criterion on the most difficult stage (stimulus duration of 2.5 s) and were therefore able to complete training, five male *Fmr1-^Δ^KH1^−/y^*, one female *Fmr1-^Δ^KH1^+/−^*, and two female *Fmr1-^Δ^KH1^−/−^* rats were unable to complete training. When we analyzed male and female WT rats compared to male *Fmr1-^Δ^KH1^+/y^* and female *Fmr1-^Δ^KH1^−/−^* rats, there was a significant association between genotype and completion of training (**Figure 3B**; 2 × 2 contingency table for genotype x completion of training, *Phi* = 0.038). There was no association between sex and completion of training (*Phi* = 0.599). Furthermore, there was no effect of sex on performance across training in WT and *Fmr1-^Δ^KH1* rats (**Supplementary Table 1**), suggesting that, much like patients with FXS, both male and female *Fmr1-^Δ^KH1* rats present with attentional deficits. Therefore, data from both sexes were pooled for visual representation and presented in two groups: WT and *Fmr1-^Δ^KH1*.

By examining the performance of all rats that completed training, we found that *Fmr1-^Δ^KH1* rats took more sessions to reach criterion in the final two stages (where the stimulus durations were 5 s and 2.5 s), compared to WT littermates (**Figure 3C**; linear mixed-effects model (LMM), for genotype x schedule across training *p* = 0.001, for genotype at 5 s, *p* = 0.003, 2.5 s, *p* = 0.015). We also found that the decline in performance was paralleled by an increase in omission rates (**Figure 3D**; LMM for genotype across training, *p* = 0.001), where the *Fmr1-^Δ^KH1* rats omitted more often than WT littermates at stimulus durations shorter than 30 s (**Figure 3D**; LMM for genotype at 20 s, *p* = 0.038, 10 s, *p* = 0.011, 5 s, *p* = 0.02, 2.5 s, *p* = 0.001). When we looked closer at these omitted responses, we found that some of them were in fact correct responses that were performed after the time allotted, which we termed “late responses”. We also found that the *Fmr1-^Δ^KH1* rats performed more of these late responses than WT littermates (**Figure 3E**; LMM for genotype across training, *p* = 0.009) at the shortest stimulus durations (**Figure 3E**; LMM for genotype at 5 s, *p* = 0.012, 2.5 s, *p* = 0.018). When the *Fmr1-^Δ^KH1* rats did make the correct choice in the time allotted, they took longer to respond than WT littermates (**Figure 3F**; LMM for genotype across training, p = 2.01 × 10^−33^, at 30 s, *p* = 0.003, 20 s, *p* = 0.009, 10 s, *p* = 0.026, 5 s, *p* = 0.032). Altogether, these findings indicate impaired sustained attention in male and female *Fmr1-^Δ^KH1* rats across training.

Importantly, these deficits were not attributed to impairments in learning or sensory perception because (1) accuracy remained unaffected across all training sessions for each stimulus duration (**Figure 3G**; LMM for stimulus duration x genotype, *p* = 0.463, and genotype, *p* = 0.924) and (2) the increased omission rates only appeared at the 20 s stimulus duration and onward. Furthermore, the deficits were not due to decreased motivation for food or motor deficits because the latency of *Fmr1-^Δ^KH1* rats to collect reward after a correct response was comparable to WT rats (**Supplementary Figure 6A**; LMM for stimulus duration x genotype, *p* = 0.671, and genotype, *p* = 0.931) and the average total amount of trials initiated and completed did not differ by genotype (**Supplementary Figure 6B**; LMM for stimulus duration x genotype, *p* = 0.755, and genotype, *p* = 0.385). Additionally, the rate of premature responses in *Fmr1-^Δ^KH1* rats was equal to WT littermates and both were low overall, suggesting that impulsivity was not a factor in the delay in reaching criterion (**Supplementary Figure 6C**; LMM for stimulus duration x genotype, *p* = 0.926, and genotype, *p* = 0.613). In summary, these results indicate that male and female *Fmr1-^Δ^KH1* rats have impairments that are specific to sustained attention, which is also commonly disrupted in FXS patients [53].

Though we did not find sex differences amongst *Fmr1-^Δ^KH1* rats, males, regardless of genotype, took longer to collect reward than females (**Supplementary Figure 7A**; LMM for sex x schedule, *p* = 0.024, for sex at 30 s, *p* = 0.022, 20 s, *p* = 0.01, 10 s, *p* = 0.005, 5 s, *p* = 0.024). Also, females, regardless of genotype, made more perseverative responses after a correct choice at 60, 30, 5, and 2.5 s (**Supplementary Figure 7B**; LMM for sex, *p* = 0.001, at 60 s, *p* = 0.012, 30 s, *p* = 0.021, 5 s, *p* = 0.023, 2.5 s, *p* = 0.004). To our knowledge, this is the first study to report sex differences in perseverative responses and reward collection latency in rats on the 5-CSRTT. Both might be explained by an overall decrease in activity in males, which are also less active in the open field test [54].

In our second analysis, we found that female *Fmr1-^Δ^KH1^+/−^* rats, which have variable expression of the FMRP with the in-frame deletion due to random X chromosome inactivation, did not have significant deficits in any of these measurements compared to their WT and *Fmr1-^Δ^KH1^−/−^* littermates (**Supplementary Figure 8**).

### Changes to the mPFC gene expression profile in the *Fmr1-^Δ^KH1* rat model of FXS

To identify gene expression differences associated with KH1 domain deletion in a brain region largely responsible for sustained attention, we applied transcriptome-wide RNA-seq and measured global gene expression profiles in mPFC samples of *Fmr1-^Δ^KH1^−/−^* and WT rats following the analytic pipeline and data pre-processing described in **Supplementary Figures 9 and 10** (see Methods). We first confirmed, based on the RNAseq data and using the Integrated Genome Viewer and the Integrated Genome Browser, that the mutation in *Fmr1* leads to exon 8 skipping (**Supplementary Figure 11**), which is in fact not detected in *Fmr1-^Δ^KH1^−/−^* rats (**Figure 4A**). Similarly, we found that the levels of RNA transcripts aligning to *Fmr1* exon sequences (except for exon 8) are comparable between WT and *Fmr1-^Δ^KH1^−/y^* samples, indicating that the decreased levels of the *Fmrp-^Δ^KH1* protein, observed in our immunoblotting analysis, is not due to reduced mRNA levels. Subsequently, we sought to identify differential gene expression (DGE) signatures and found 259 up- and 297 down-regulated genes in *Fmr1-^Δ^KH1^−/y^* rats compared to WT (using False Discovery Rate (FDR) < 0.1) (**Figure 4B, Supplementary Table 2**). Notably, these genes were mainly associated with differences in genotype and not with any other factor, including differences in parents, RIN values, age, date of dissection or estimated cell type proportions (**Supplementary Figure 10H**). Consistent with this result, DGE signatures largely separated the two genotypes (**Figure 4C**).

**Figure 4.**
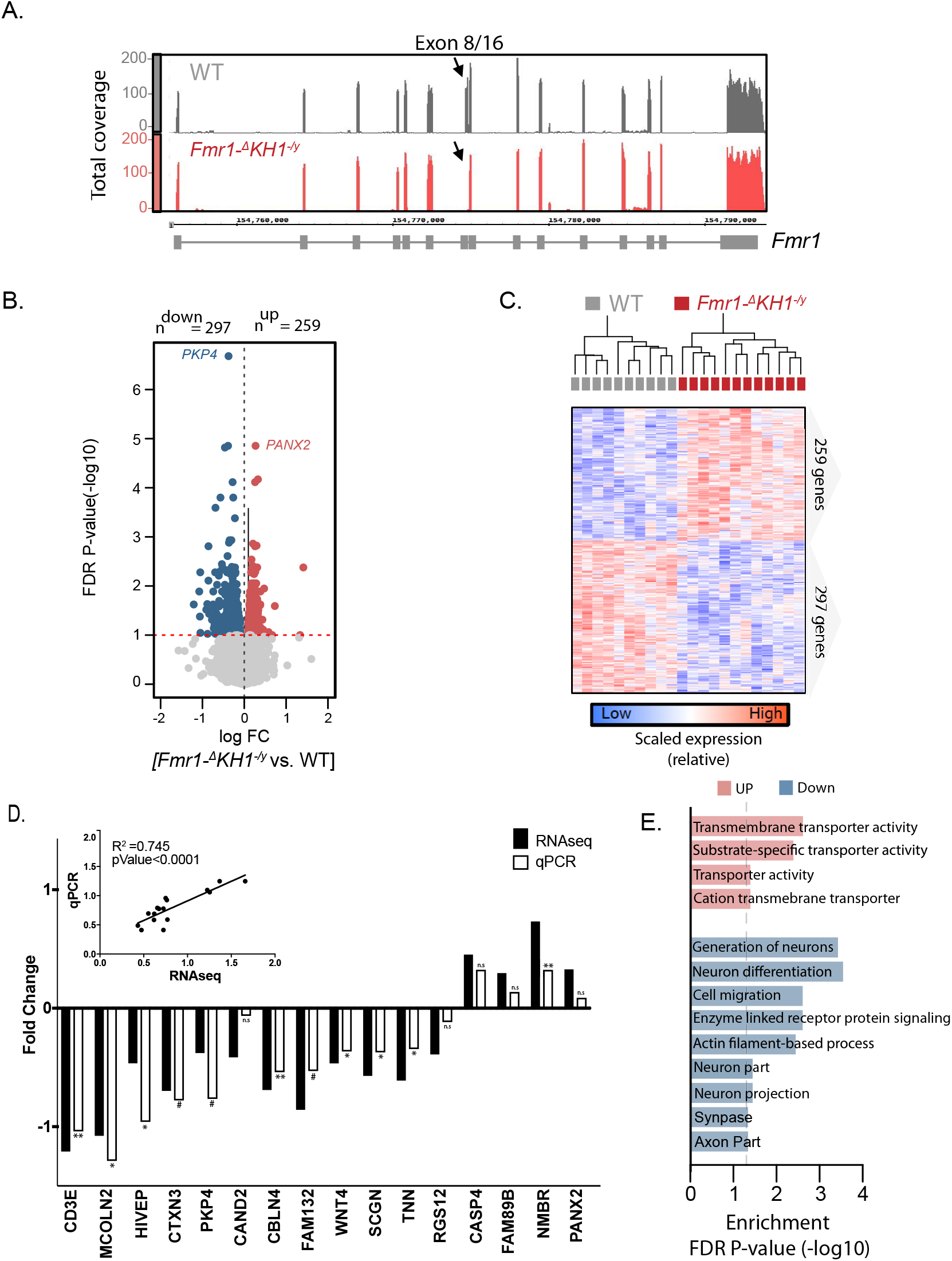
Comparative RNA-seq analysis between *Fmr1-^Δ^KH1^−/y^* and WT rats. (**A**) Depth of RNA-seq coverage (y-axis) plots across all 16 exons of the *FMR1* gene (x-axis) using the Integrative Genome Browser (IGB) viewer. Plots represent pooled coverage across all animals for each genotype (WT=10, *Fmr1-^Δ^KH1^−/y^* =12). (**B**) Volcano plot comparing the extent of FDR *q*-value significance (y-axis) and log fold change (x-axis) for differential gene expression (DGE) between *Fmr1-^Δ^KH1^−/y^* and WT rats. (**C**) DGE signatures segregate *Fmr1-^Δ^KH1^−/y^* and WT samples. Normalized editing levels (z-scores) were used in hierarchical clustering. Each row corresponds to one gene and each column corresponds to one sample. (**D**) qPCR validation on 16 genes, identified to be differentially expressed by our RNAseq analysis. Validation was done on an independent set of *Fmr1-^Δ^KH1^−/y^* rats and WT littermate mPFC samples (n=7/genotype). 8 genes showed statistically significant changes (one tail t-test, *p*<0.05) and 3 showed a trend towards significance (one tail t-test, *p*<0.1). Inset shows the significant correlation between the absolute values of the RNAseq and qPCR mean fold changes. (**E**) Functional annotation of DGE signatures, parsed by up- and down-regulated genes.

To validate our findings in an independent cohort of rats (n=7/genotype), we used quantitative reverse transcription PCR (RT-qPCR) on a cross section of the DEG, including both up and down regulated genes, for a total of sixteen DEGs. Analysis of the correlation between the two studies (i.e. absolute values of the RNAseq and qPCR mean fold changes) showed a significant correlation (*R*^2^= 0.74, *p*<0.0001) (**Figure 4D**). Moreover, eight out of the sixteen DEGs showed statistically significant changes (one tail *t*-test, *p*<0.05) and three showed a trend towards significance (one tail *t*-test, *p*<0.1) (**Figure 4D**).

Next, to gain biological insights into the function of the differentially expressed genes, we performed gene ontology enrichment analysis (**Figure 4E**). We found that up-regulated genes were enriched in biological processes that included transmembrane transporter activity (q-value FDR B&H <4.02×10^−02^). In parallel, down-regulated genes were enriched for (1) cellular components, including neuron part (q-value = 2.45×10^−02^), neuron projection (q-value = 2.45E-02), synapse (q-value = 2.85×10^−02^) and axon part (q-value = 3.74×10^−02^) and (2) biological processes including generation, differentiation, and migration of neurons (q-value = 4.57×10^−02^), enzyme linked receptor protein signaling pathway (q-value = 4.57×10^−02^), and actin filament based processes (q-value = 4.57×10^−02^) (**Supplementary Table 3**).

We then tested whether the *Fmr1-^Δ^KH1* DGE signatures were enriched for Fmrp targets [2] or risk genes for ASD [55–57], ID [58], or genes found with *de novo* mutations in schizophrenia (SCZ) [59]. We found that the down-regulated genes in *Fmr1-^Δ^KH1^−/y^* rats show significant overlap with Fmrp targets (∩=33, FDR *P*=0.0002) and SCZ genes (∩=7, FDR *P*=0.03) (**Supplementary Figure 12A**).

### Rat mPFC gene networks, preserved in human frontal cortex, are altered by the loss of the Fmrp-KH1 domain

Next, we asked whether mPFC gene networks in WT rats are conserved in human and if any of the conserved networks were especially vulnerable to the effects of *Fmr1-^Δ^KH1* deletion. To address these questions, we first built a reference WT co-expression network by combining mPFC RNA-seq data across 35 WT rats, matched for age and sex, and using weighted gene co-expression network analysis (WGCNA) (see Methods). Our WGCNA analysis identified 23 modules specific to the mPFC of WT rats (**Supplementary Table 2**). Next, we determined whether the co-expression patterns of these 23 modules were preserved in the human brain. For this purpose, we created separate transcriptional networks from previously published human cortex tissue (BA 9/41) sampled from control individuals [60] in order to systematically explore potential species similarities and differences. Inter-species co-expression preservation has been shown to prioritize disease gene selection under genetic disease loci [61, 62] and to categorize the function of poorly characterized genes better than co-expression in a single species. This approach is sensitive to detecting fundamental differences in the underlying gene co-regulatory patterns between WT rats and unaffected humans, and vice versa, as being preserved or disrupted. Using a permutation-based preservation statistic (*Z_summary_*) with 500 random permutations, we observed strong to moderate preservation between the two species (all network modules displayed a *Z_summary_* score >2, which was higher than a random sampling of 100 genes), indicating similar levels of gene co-regulation between rat mPFC and human frontal cortex (**Figure 5A**).

**Figure 5.**
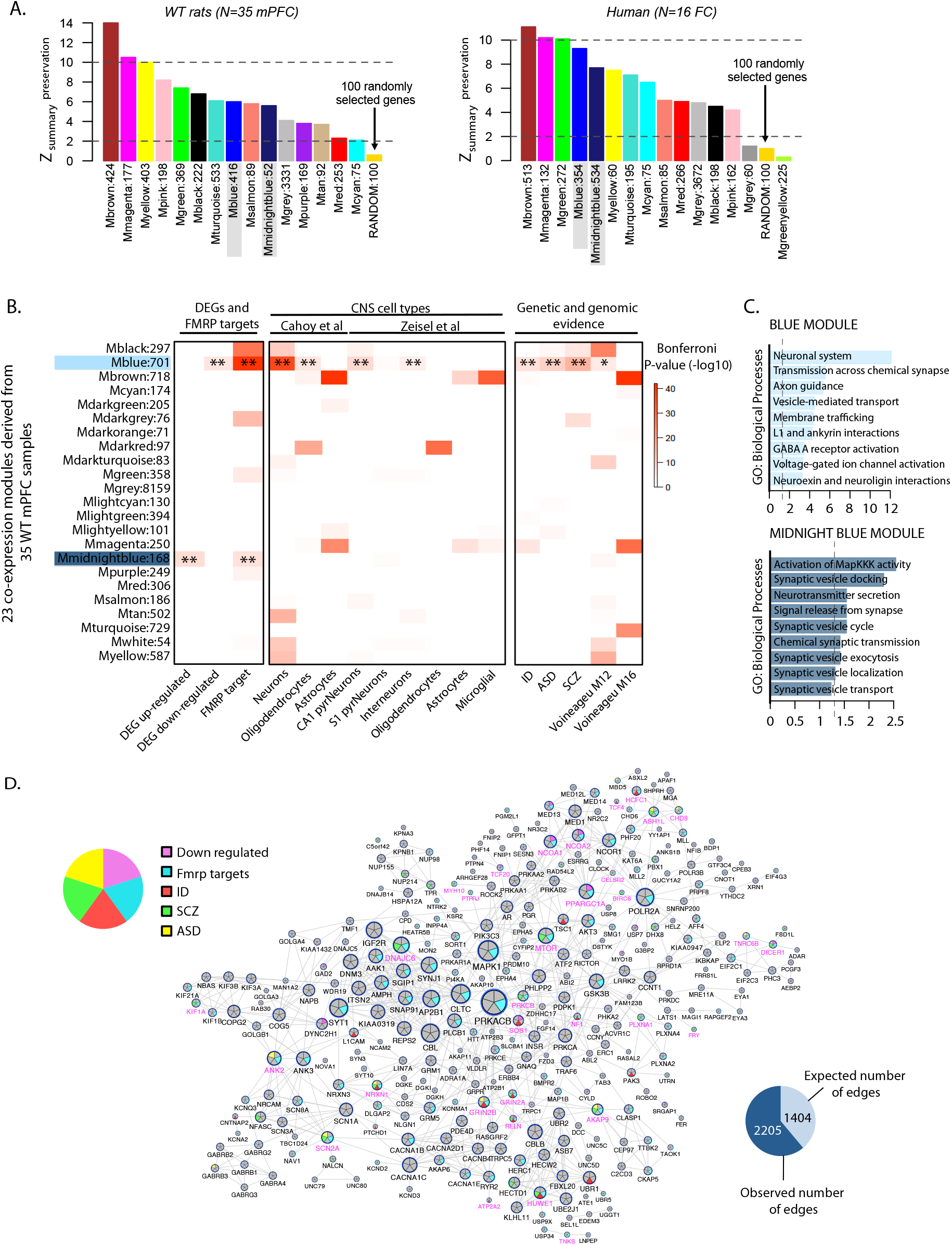
mPFC gene co-expression networks of WT rats and preservation with human frontal cortex co-expression networks. WGCNA identified 23 modules across 35 WT mPFC samples. (**A**) Permutation-based preservation statistic (*Z_summary_*) (y-axis) were generated with 500 random permutations across all WT modules (x-axis), comparing WT rat mPFC (left) and healthy human frontal cortex (right) preservation to 100 randomly selected genes. *Z_summary_*<2 indicate no evidence of preservation, *2<Z_summary_*<10 implies moderate preservation, and *Z_summary_* >10 suggests strong preservation. These results indicate that the blue and midnightblue modules, which are significantly enriched for DEGs, are indeed preserved. (**B**) All modules were assessed for enrichment of *Fmr1-^Δ^KH1* DGE signatures and Fmrp targets, CNS cell type specific signatures, genetic risk loci and genomic evidence of neurodevelopmental disorders. Overrepresentation analysis of these gene sets within DGE signatures and WT transcriptome modules was analyzed using a one-sided Fishers exact test to assess the statistical significance. All P-values, from all gene sets and modules, were adjusted for multiple testing using Bonferroni procedure. (**C**) Functional annotation of the blue (top) and midnightblue (bottom) modules. (**D**) Protein interaction network for blue module genes shows a significant overrepresentation of high confidence direct protein interactions, beyond what was expected by chance (*P*<0.0001). Hub genes include numerous Fmrp target genes and several ID, schizophrenia (SCZ), and ASD genetic risk loci. Genes in pink lettering were differentially expressed in *Fmr1-^Δ^KH1^−/y^* rats.

To assess whether WT rat modules were vulnerable to the *Fmr1-^Δ^KH1* deletion, we tested for enrichment of the *Fmr1-^Δ^KH1^−/y^* DEGs. Of the 23 identified modules, one module (blue) contained a strong, significant over-representation for *Fmr1-^Δ^KH1^−/y^* down-regulated genes and FMRP targets, and another module (midnightblue) that was enriched for *Fmr1-^Δ^KH1* up-regulated genes and FMRP targets (**Figure 5B**). The blue module also contained a significant enrichment for neuronal cell type markers and genes implicated in ASD, ID and SCZ (**Figure 5B, Supplementary Table 2**). Functional annotation of the blue module revealed enrichment primarily associated with synaptic signaling, gated channel activity and neuronal system-related terms (**Figure 5C**). The midnightblue module did not display any cell type specificity nor any enrichment for risk genes. Functional annotation of the midnightblue module revealed functional terms implicating MapKKK activity, synaptic vesicle docking and neurotransmitter secretion (**Figure 5C**).

Subsequently, we tested whether genes that are co-expressed together in the blue module indeed interact with each other at the protein level. A significant overrepresentation of high-confidence direct protein interactions was identified in the blue module, beyond what was expected by chance (*P*<0.0001) (**Figure 5D**). Hub genes in the blue module include numerous FMRP target genes including *MTOR, ANK2, ANK3, SCN2A, GRIN2A, RELN*, and *NRXN1* [2]. Interestingly, all of these genes are implicated in neurodevelopmental disorders. *ANK2, SCN2A, and NRXN1* are top risk genes for ASD [57], and the others are associated with neurodevelopmental syndromes (e.g., *MTOR* in Smith-Kingsmore syndrome (MIM 616638), *ANK3* in an autosomal recessive ID syndrome (MIM 615493), *GRIN2A* in a form of focal epilepsy and speech disorder with or without ID (MIM 245570), *RELN* in a lissencephaly syndrome (MIM 257320), and *NRXN1* in Pitt-Hopkins-like syndrome (MIM 614325). This module also includes the *TSC1* gene, which is associated with Tuberous Sclerosis (MIM 191100).

We also observed another module (midnightblue) that was enriched for *Fmr1-^Δ^KH1* up-regulated genes and FMRP targets (**Figure 5B**). This midnightblue module did not display any cell type specificity nor any enrichment for risk genes. Functional annotation of the midnightblue module revealed functional terms implicating MapKKK activity, synaptic vesicle docking and neurotransmitter secretion (**Figure 5C**).

## Discussion

This study is the first to uncover that the model published as a *Fmr1* KO rat is not a null KO of *Fmr1*, but rather a model for the deletion of the KH1, an FMRP domain that is responsible for RNA-binding [63]. Further, we find that deletion of KH1 causes attentional impairments that parallel phenotypes seen in FXS patients and *Fmr1* KO mice and leads to alterations in the transcriptional profiles within the mPFC, which are of potential translational value for subjects with FXS.

These results, and prior results with this rat model, indicate that the specific role of the KH1 domain within Fmrp is vital for many of the molecular, morphological, and functional phenotypes observed in FXS models that are often attributed to a loss of Fmrp. However, the specific role of the KH1 domain in these functions is understudied. In an individual where a point mutation in KH1 led to FXS, the mutant FMRP was shown to be unable to bind known mRNA targets of Fmrp, associate to polyribosomes, and traffic AMPA receptors in a mGluR-mediated manner [14]. In this individual, disrupted function of the KH1 domain was sufficient to cause the classic symptoms of FXS that usually follow from silencing of the entire *FMR1* gene, including attention deficits.

Cognitive deficits observed in FXS rodent models are often subtle, specific to one strain or species, or not detected [34, 47, 64–68]. We believe that this is partially due to the fact that the focus of these studies has been on the hippocampus as the locus of cognitive dysfunction. For example, deficits in spatial memory on the reversal phase of the Morris Water Maze, which is hippocampal-dependent, have been reported as minor in the *Fmr1* KO mouse [64] and have not been recapitulated in the *Fmr1-^Δ^KH1* rat [34] in the Morris Water Maze or, in this study, in a related task, the Barnes maze. Instead, we found that *Fmr1-^Δ^KH1* rats were impaired in acquiring an mPFC-dependent attentional task, the 5-CSRT task, to standard performance criterion at a 5 s stimulus duration or less. The delayed performance was due to a deficit in sustained attention, exemplified by increased response latency and omissions. This deficit resembles the impairments seen in individuals diagnosed with FXS [69, 70] and *Fmr1* KO mice [71]. Notably, Sprague Dawley rats typically can perform the task at 1 s or less [52, 72], while *Fmr1-^Δ^KH1* rats were not able to perform the task at stimulus durations shorter than 2.5 s, suggesting a relatively severe deficit in sustained attention. We recently reported a similar deficit in sustained attention in a SΛank3-deficient rat model of ASD [52], suggesting convergent findings.

The basis of this deficit in sustained attention could be attributed to functional impairments in the mPFC of *Fmr1-^Δ^KH1* rats. Dysregulated sustained attention has been shown to follow manipulations of mPFC activity in rats that have previously acquired the task. Lister hooded rats that underwent treatment with an immunotoxin to deplete cholinergic function in the nucleus basalis magnocellularis (nbm) of the basal forebrain, which sends cholinergic projections to the medial frontal cortex, had increased omissions and no difference in accuracy [73]. Increased omissions and response latency were also observed in rats with lesions to mPFC or imbalanced inhibition/disinhibition in mPFC [35, 74]. The performance of *Fmr1-^Δ^KH1* rats during the acquisition of the task mirrors the performance of WT rats with specific manipulations of mPFC activity, indicating that the mPFC is implicated in the manifestation of these attentional deficits in the *Fmr1-^Δ^KH1* rat model and could be a result of an insult to the mPFC by the *Fmr1* mutation during early developmental stages. *Fmr1* KO mice also had deficits in the acquisition of the 5-CSRT task, which was accompanied by alterations in prefrontal synaptic composition and neural activity [30]. Therefore, the mPFC warrants further study as the basis of cognitive impairment in this *Fmr1-^Δ^KH1* rat model of FXS.

Our approach to address this was to probe the molecular profile of the mPFC following the loss of the KH1 domain using RNAseq analysis. We observed hundreds of dysregulated genes (FDR 10%) associated with *Fmr1-^Δ^KH1*. Up-regulated genes were enriched in biological processes that included transmembrane transporter activity. Genes within this GO category included several solute carrier proteins including a member of the Na^+^/H^+^ exchanger (NHE) superfamily, *SLC9A9*. This family of exchangers controls ion transport across membranes, which is essential for regulating cellular pH and electrical excitability that is known to be affected in FXS [75]. SLC9A9 is highly expressed in the brain and mutations in the encoding gene have been associated with ASD [76], ADHD [77–79], and epilepsy, which are all prevalent in FXS [75].

Down-regulated genes were enriched for neural and synaptic components and for biological processes including generation, differentiation, and migration of neurons and actin filament based processes. Impaired actin cytoskeletal function has consistently been reported in FXS models [80] and is thought to underlie the abnormal dendritic spine phenotype common to subjects with FXS [80]. Pkp4 is a member of a subfamily of armadillo proteins known to regulate cell adhesion and cytoskeletal organization [81, 82] and is a validated target of Fmrp [2, 81, 83]. Notably, our transcriptional analyses show that Pkp4 mRNA is down-regulated in *Fmr1-^Δ^KH1* rats and is the highest differentially expressed gene in our analysis. Down regulation of Pkp4 mRNA and other Fmrp targets at the transcriptional level could be an indirect consequence of the Fmrp mutation and an attempt by the neurons to compensate for the increased translation of Fmrp targets caused by the lack of translational inhibition by Fmrp.

Results derived from our reference WT transcriptome co-expression network echo our DEG findings and further refine the biological processes involved in *Fmr1-^Δ^KH1*. Functional annotation of the blue module, which we found to be conserved in human PFC co-expression networks and to be significantly enriched for down-regulated *Fmr1-^Δ^KH1-related* genes, known Fmrp targets, neuronal cell type signatures, and genes implicated in ID, ASD and SCZ, revealed enriched GO terms including neuronal system, axon guidance, and neurexin (an Fmrp target [2]) and neuroligin interactions. These biological processes were previously reported to be dysregulated in a transcriptomic study of the cerebellum of *Fmr1* KO mice [84] and in functional studies of Fmrp [85, 86]. Amongst the hub genes in the blue module are numerous FMRP target genes and several ASD risk genes, including: *MTOR, ANK2, ANK3, TSC1, SCN2A, GRIN2A, RELN* and *NRXN1*. The midnightblue module was enriched for up-regulated genes and Fmrp targets and revealed functional terms implicating MapKKK activity, synaptic vesicle docking and neurotransmitter secretion.

In summary, we have shown here that a specific deletion of the KH1 domain is sufficient to cause FXS-like phenotypes in rat. The behavioral task we employed provides a tool to screen potential therapeutic candidates for efficacy in treating a highly common cognitive deficit in these rats that is seen in both males and females diagnosed with FXS, dysregulated attention, which is associated with mPFC dysfunction. In addition, the results from our RNAseq analysis of the mPFC supply multiple potential treatment avenues to explore. Now that these deficits are elucidated in the *Fmr1-^Δ^KH1* rat, we can begin to uncover their underlying circuit mechanisms by probing the mPFC with *in vivo* imaging and electrophysiology.

## Materials and Methods

### Generation of the *Fmr1-^Δ^KH1* rat model

The *Fmr1-^Δ^KH1* rat model, previously reported as the *Fmr1* KO rat model [33, 37–40], was generated using zinc-finger nucleases (ZFN) in the outbred Sprague-Dawley background. The design and cloning of the ZFN, as well as the embryonic microinjection and screening for positive founder rats were performed by SAGE Labs (Boyertown, PA USA) as previously described [87]. The best performing ZFN pair targeting the CATGAACAGTTTATCgtacgaGAAGATCTGATGGGT sequence, located between 18631bp–18666bp in the *Fmr1* gene (NCBI reference sequence NC_005120), were used for embryo microinjection. Positive Sprague-Dawley founder animals with a deletion in the *Fmr1* gene were mated to produce F1 breeder pairs. PCR amplification at the target sites followed by sequencing analysis revealed the exact deletion of 122bp at the junction of intron 7 and exon 8 (between 18533bp-18654bp), as previously described by Hamilton et al. [33].

### Animal breeding, care, and husbandry

This study used age-matched male and female littermate rats. To produce WT and *Fmr1-^Δ^KH1* hemizygous (*Fmr1-^Δ^KH1^−/y^*) male littermates we bred heterozygous (*Fmr1-^Δ^KH1^+/−^*) females with WT males. To produce WT, *Fmr1-^Δ^KH1^+/−^* and *Fmr1-^Δ^KH1^−/−^* female littermates we bred *Fmr1-^Δ^KH1^+/−^* females with WT and *Fmr1-^Δ^KH1^−/y^* males. WT and hemizygous males were used for the both RNAseq analyses and the attentional task and WT, heterozygous, and homozygous females were used for the attentional task. For the attentional task, WT rats were housed with *Fmr1-^Δ^KH1^+/−^, Fmr1-^Δ^KH1^−/−^*, or *Fmr1-^Δ^KH1^−/y^* sex-matched littermates. All rats were kept under veterinary supervision in a 12 h reverse light/dark cycle at 22±2°C. Animals were pair-caged with food and water available *ad libitum*. Rats tested on the 5-CSRTT were food restricted to 85% of their free-feeding weight. All animal procedures were approved by the Institutional Animal Care and Use Committees at the Icahn School of Medicine at Mount Sinai.

### Testes weight

Testes were dissected from 10-week-old male *Fmr1-^Δ^KH1^−/y^* rats (n=19) and WT littermates (n=19). After gonadal fat pads were removed and testes were weighed, testes:body weight ratios were calculated. The data was analyzed with a two-tailed T-test.

### Total lysate preparation

mPFC tissues were dissected from 8-weeks old rats as previously described [88]. Tissues were homogenized in 100ul ice cold RIPA buffer supplemented with 1:100 proteinase inhibitors cocktail (Thermo Scientific) and 1:100 phosphatase inhibitors (Thermo Scientific), using a Teflon-glass homogenizer. The homogenate was centrifuged at 12000g for 20 min at 4°C. The recovered supernatant was centrifuged again at 12000 rpm for 20 min at 4°C. The protein concentration in the final recovered supernatant was determined using the BCA protein assay (Pierce).

### Structural data

FMR1 orthologs from 58 species, including *D. melanogaster*, two Enterogona (Chordata: Tunicata), 12 fishes, *X. tropicalis*, two reptiles, five birds and 35 mammals, were extracted from Ensembl and aligned using Alvis v. 0.1 software. The X-ray structure of the human FMRP KH1-KH2 domains (PDBID = 2QND) was generated using Pymol v1.7.2.1.

### Immunoblotting

Immunoblotting was performed using a standard protocol [89]. Briefly, 10 μg of each protein lysate were loaded to a 4-12% SDS-polyacrylamide gel electrophoresis (PAGE gel, Invitrogen; Carlsbad, CA USA), which was transferred to polyvinylidene fluoride membrane for immunoblotting. For Fmrp detection we used the anti-Fmrp (G468) antibodies targeted against the C-terminus of the Fmrp (1:1000, Cell Signaling) and the anti-Fmrp (F3930) antibodies targeted against the N-terminus of Fmrp (1:1000, Sigma Aldrich). The Anti beta III tubulin antibodies (1:2000, Abcam; ab18207, RRID: AB_444319) were used to quantify the beta III tubulin level, used for normalization. HRP-conjugated anti-rabbit (1:5,000), BRID: AB_2337910 HRP-conjugated anti-mouse antibodies (1:5,000), BRID: AB_2340031 were purchased from Jackson ImmunoResearch Laboratories (West Grove, PA USA). ECL substrate (Pierce; Thermo Scientific, Rockford, IL USA) or SuperSignal West Femto (Thermo Scientific, Rockford, IL USA) substrates were used to produce the signal that was detected on a G:Box Chemi-XT4 GENESys imager (Syngene; Cambridge UK). Blots were quantified using the software GeneTools (SynGene, version 4.02).

### Immunoprecipitation followed by Fmrp-^Δ^KH1 Protein sequencing

Frontal cortex were dissected from 8-week old WT and *Fmr1-^Δ^KH^−/y^* rats and homogenized in lysis buffer (100 mM NaCl, 10mM MgCl_2_, 10 mM Tris-HCl [90], 1 mM diothiolhreitol, 1% Triton X-100, (1:100) proteinase inhibitors cocktail (Thermo Scientific) and (1:100) phosphatase inhibitors (Thermo Scientific), using an electric tissue homogenizer [91]. Samples were incubated on ice for 5 minutes and centrifuged at 12,000 x g for 8 min at 4°C. The recovered supernatant was centrifuged again at 12,000 × g for 8 min at 4°C. The protein concentration in the final recovered supernatant was determined using the BCA protein assay (Pierce). 800 ug of protein extract was used for Fmrp immunoprecipitation experiments. Fmrp was immunoprecipated based on the previously established protocol [83]. Briefly, Fmrp was immunoprecipitated with 6.24 ug 7G1-1 *Fmr1* monoclonal mAb conjugated to 1.5 mg of Protein A Dynabeads (Invitrogen). The same amount of monoclonal mouse IgG2B (R&D Systems) was used as control. The immunoprecipitates as well as 20 ug of frontal cortex protein lysate from WT and *Fmr1-^Δ^KH1^−/y^* rats were loaded onto 4-12% SDS-polyacrylamide gel electrophoresis (PAGE gel, Invitrogen; Carlsbad, CA USA) and ran for 3 hours at 200 V, 60 mA to allow an optimal separation of the bands around 75 MW followed by transfer onto PVDF membranes (Invitrogen) using XCell II Blot Module system (Thermo Scientific). Membranes were immunoblotted with anti-Fmr1 Abs (1:1000, N-Terminal, Sigma Aldrich F3930-25UL). Subsequently, membranes were incubated with appropriate anti-rabbit HRP conjugated secondary antibodies (1:5000, Jackson ImmunoResearch Laboratories). Images were developed using the enhanced chemiluminescent substrate West Femto (Thermo Scientific).

### PCR amplification on genomic DNA followed by Sanger sequencing

Tail samples were collected from WT and *Fmr1-^Δ^KH1^−/y^* rats and DNA was extracted using the QIAamp DNA Mini Kit (Qiagen) according to the manufacturer’s instructions. 25ng of each samples was PCR amplified using the Fmr1-G-F and Fmr1-G-R primers. PCR products were loaded on agarose gel and pure bands from each of the WT and *Fmr1-^Δ^KH1* samples were sliced from the gel and cleaned using the QIAquick Gel Extraction Kit (Qiagen). Purified samples were sent to GENEWIZ for Sanger sequencing using both, the Fmr1-G-F and Fmr1-G-R primers. Primers location and sequence are described in **Supplementary Table 4**.

### Reverse transcriptase PCR (RT PCR) followed by Sanger sequencing

mPFC tissues were dissected from 8-weeks old WT and *Fmr1-^Δ^KH1^−/y^* male rats as previously described [88]. RNA was extracted using the RNeasy Mini Kit (Qiagen) according to the manufacturer’s instructions. 2ug of RNA was then used to prepare cDNA, using the SuperScript II Reverse Transcriptase Kit (Invitrogen) and following the manufacture’s instructions. 100 ng of each sample was PCR amplified using the F-Fmr1, which aligns to exons 6/7 junction and the R-Fmr1, which aligns to exons 11/12 junction. PCR products were loaded on agarose gel and pure bands from each of the WT and *Fmr1-^Δ^KH1* samples were sliced from the gel and cleaned using the QIAquick Gel Extraction Kit (Qiagen). Purified samples were sent to GENEWIZ for Sanger sequencing using both, the F-Fmr1 and R-Fmr1 primers. Primers location and sequence are described in **Supplementary Table 4**.

### Open field test

Rats were exposed to a brightly lit novel 90 cm × 90 cm environment during their light-cycle for one hour. All horizontal movements were automatically tracked by Noldus Ethovision system and samples were analyzed in 10-minute bins. Grooming, jumping, and rearing were scored manually. After a significant effect of time and no significant effect of genotype were discovered in a first group of animals (WT: n=6, *Fmr1-^Δ^KH1^−/y^:* n=7), the experiment was repeated and the same effect was replicated in a second group (WT: n=7, *Fmr1-^Δ^KH1^−/y^:* n=5). The data was analyzed with SPSS statistical package, version 23 (IBM SPSS Statistics, Armonk, North Castle, NY, USA) using repeated measures ANOVA where time was the within-subjects factors, genotype was the between-subjects factor, and batch was a covariate.

### Barnes maze

In this assay, similar to what has been conducted before [92], rats were trained to navigate around a brightly lit circular arena with 18 holes around the edge to find the hole with an escape box (a box full of bedding) underneath it using spatial cues. Once the rat found the escape box, the trial ended. If the rat did not find the hole after three minutes, it was guided to it. Male WT (n=6) and *Fmr1-^Δ^KH1^−/y^* (n=7) rats were trained four times a day, starting each trial in the middle of the area facing north, for four days in a row. The next day, they were given a probe test where no escape box was present and their time in the quadrant with the target hole, the one that used to contain the escape box, was measured. Subsequent to the probe test, they were trained on the reversal phase of the task, where the escape box was instead placed under a hole 180° away from the original target goal, for three days, followed by another probe test. Two weeks later, a final probe test was conducted in order to assess long-term memory. Data was analyzed with SPSS statistical package, version 23 (IBM SPSS Statistics, Armonk, North Castle, NY, USA) and included repeated measures ANOVA where day was the within-subjects factor and genotype was the between-subjects factor.

### 5-CSRTT

The 5-CSRTT was carried out as we have previously described [52], with slight modifications due to performance. Briefly, training on the 5-CSRTT began when the rats were 8-weeks old and after they were habituated to being handled and food deprived to achieve ~85% of free feeding weight. Rats were first trained to touch the location of an illuminated white square presented at 1 of 5 locations on a Bussey-Saksida capacitive touchscreen system (Lafayette Instrument Company; Lafayette, IN USA) using ABET II Software for Touch Screens. If a capacitive screen touch occurred at the illuminated location during the stimulus presentation or during the subsequent limited hold period, sucrose (valve open for 250 ms) was delivered in the reward receptacle located across the chamber from the touch screen. Training occurred in stages, where the light stimulus duration decreased from 60 to 30 to 20 to 10 to 5 to 2.5 s. The limited hold period lasted for the duration of the stimulus presentation at stimulus durations of 60 s to 5 s and lasted for 2.5 s after a stimulus duration of 2.5 s. Rats advanced schedules once criterion performance was met. Criteria for progression were an accuracy rate higher than 80% (accuracy rate = correct trials/total trials) and an omission rate lower than 20% (omission rate = omitted trials/total trials). Trials where the rat made a correct response after the limited hold period were termed “late responses.” Once criterion was reached with a stimulus duration of 2.5 s, training was recorded as complete. Four separate batches were trained on the 5-CSRTT, totaling 12 WT males, 15 *Fmr1-^Δ^KH1^−/y^* males, 11 WT females, 13 *Fmr1-^Δ^KH1^+/−^* females, and 12 *Fmr1-^Δ^KH1^−/−^* females rats, where the experimenter was blind to subject genotype. Five *Fmr1-^Δ^KH1^−/y^* males, one *Fmr1-^Δ^KH1^+/+^* female, one *Fmr1-^Δ^KH1^+/−^* female, and two *Fmr1-^Δ^KH1^−/−^* females that either did not reach criterion on a stimulus duration of 5 s after 30 sessions or on a stimulus duration of 2.5 s after 45 sessions were removed from analysis. When subsets of 10 animals per group were randomly sampled from the male and female WT and *Fmr1-^Δ^KH1* dataset 10,000 times and analyzed via a two-way ANOVA (genotype × sex), a significant main effect of genotype (*p*<0.05) on the % omissions measure at a 2.5 s stimulus duration was observed 92% of the time, suggesting that there was reproducibility of this finding regardless of batch. Therefore, we focused our analysis on the pooled data. Because *Fmr1* is X-linked, there was an imbalance in the number of genotypes available between males (two) and females (three). Thus the analysis included two separate comparisons: WT male and female rats that were compared to *Fmr1-^Δ^KH1^−/y^* male and *Fmr1-^Δ^KH1^−/−^* female rats, and WT female rats that were compared to *Fmr1-^Δ^KH1^+/−^ and Fmr1-^Δ^KH1^−/−^* female rats. Data analysis of training data was comprised of linear mixed-effects modeling (LMM) where sex, genotype, and stimulus duration were fixed factors and rat was nested into batch as a random factor using custom scripts written in the R statistical programming environment (R Development Core Team, 2006).

### RNA isolation, library preparation and data availability

RNA was extracted from mPFC tissues from 8-weeks old WT (n=12) and *Fmr1-^Δ^KH1^−/y^* littermate (n=12) rats using the RNeasy Mini Kit (Qiagen), according to the manufacturer’s instructions. Subsequently, the quantity of all purified RNA samples was measured on a nanodrop (2.07±0.01 A_260/280_; 2.11±0.19 A_260/230_) and the quality and integrity measured with the Agilent 2100 Bioanalyzer (Agilent, Santa Clara, CA, USA). All RNA integrity numbers were greater than 9 (9.6±0.3). Following, 1μg of total RNA was used for the preparation of the RNAseq library using the Illumina Genome Analyzer IIx TruSeq mRNA Seq Kit supplied by Illumina (Cat # RS-122-2001). Poly-A-based mRNA enrichment step was carried out and cDNA was synthesized and used for library preparation using the Illumina TruSeqTM RNA sample preparation kit as previously described, except for the following steps: adapter-ligated DNA fragments were size-selected by gel-free size selection using appropriate concentration of SPRI AMPure beads to get an average 200bp peak size in adaptor ligated DNA. The size selected adaptor-ligated DNA fragments were amplified by LM-PCR. Then, Illumina recommended 6bp barcode bases were introduced at one end of the adaptors during PCR amplification step. The amplified PCR products were then purified with SPRI AMPure XP magnetic beads to get the final RNA-seq library, which was used for high-throughput RNA-seq.

All samples were sequenced on the Illumina Genome Analyzer IIx. A total of 40 million 100bp paired-end sequences were used to reliably assess expression for each sample. Overall, the design of the experiment was as follows: 12 barcoded samples/per brain region, among which 6 of each genotype, were pooled and loaded on 2 lanes, so that each sample is spread over two lanes to further minimize confounds, specifically those associated with lane effects. These raw RNA-seq fastq data have been submitted to Gene Expression Omnibus (http://www.ncbi.nlm.nih.gov/geo/) under the accession number GSEXXXXX *(accession to follow publication)*.

### Short read mapping and quantification of gene expression

All high quality short reads were mapped to the rat reference genome rn4 using the STAR Aligner v2.4.0g1 [93] with 2-pass mapping strategy (--twopassMode Basic). RNA-sequencing read quality was checked with FastQC the RNA-seqQC tools. Uniquely mapped reads overlapping genes were counted with featureCounts v1.4.4 [94] parameters: *featureCounts -T 10 -p -t exon -g gene_id)*.

### Data pre-processing

Raw count data measured 16,499 transcripts across all samples (12 WT and 12 *Fmr1-^Δ^KH1^−/y^* rats). Nonspecific filtering required more than 2 counts per million (cpm) in at least 12 samples and retrained 14,745 transcripts. Filtered raw count data was subjected to conditional quantile normalization [95] (CQN) to remove systematic bias introduced by GC-content and correct for global distortions, resulting in a normally distributed data matrix. Normalized data were inspected for outlying samples using unsupervised clustering of samples (Pearson’s correlation coefficient and average distance metric) and principal component analysis to identify outliers outside two standard deviations from these grand averages. Based on these metrics, two outliers were removed from these data (WT=2). Rattus ENSEMBL symbols were converted to HGNC symbols, then converted to human orthologues using Ensemble biomart conversion (http://www.ensembl.org/biomart). In order to form the bases for cross-species comparisons (rat, human), we constructed one large reference transcriptome of the mPFC in WT rats by integrating RNA-seq gene expression data from an additional 24 WT rats (N_tot_=35). These data were processed in an identical fashion as described above, and were generated in three batches (i.e. different processing dates). Combat batch correction [96] was applied to resolve systematic sources of variability across batches (**Supplementary Figure 13**). Finally, to estimate the relative frequencies of brain cell types for each bulk frontal cortex sample, Cibersort deconvolution analysis was applied (https://cibersort.stanford.edu/). Cibersort [97] relies on known cell subset specific marker genes and applies linear support vector regression to estimate the relative frequencies of cell types from bulk tissue. As a signature matrix, we used *a priori* defined brain cell type specific RNA-sequencing expression markers [98] to obtain estimates for neurons, oligodendrocytes and astrocytes in all frontal cortex samples.

### Visualization of the *Fmr1* deletion and splice junctions

To enable in-depth visualization of the deletion in exon 8 of the *Fmr1* gene, we used depth of coverage plots from the Integrated Genome Browser (IGB) (http://bioviz.org/igb/). The *Rattus norvegicus* reference genome version Nov_2004 was used as a reference genome, and two pooled BAM alignment files were used as input, one for each genotype (WT and *Fmr1-^Δ^KH1)*. Subsequently, to visualize predicted splice junctions in the *Fmr1* gene, we used the in Integrative Genome Viewer (IGV) (http://software.broadinstitute.org/software/igv/). Similarly, the *Rattus norvegicus* reference genome version Nov_2004 was selected, and sorted BAM files from each rat were loaded separately (10 WT and 12 *Fmr1-^Δ^KH1)* and then pooled across genotypes to show the total numbers per group.

### Differential gene expression analysis

Differential gene expression (DGE) signatures between *Fmr1-^Δ^KH1* and WT rats were identified using moderated t-tests in the *limma* package [99]. The covariates RIN, parents, age and date of dissection were included in the models to adjust for their potential confounding influence on gene expression between-group main effects and *P*-value significance was set to a FDR-corrected p-value of <0.05. This assumption was later relaxed to FDR-corrected p-value of <0.1 to yield sufficient amount of genes for down-stream network enrichment analyses.

### Weighted gene co-expression network analysis

Weighted gene co-expression network analysis [100] (WGCNA) was used to build signed coexpression networks. To construct each network, the absolute values of Pearson correlation coefficients were calculated for all possible gene pairs, and resulting values were transformed with an exponential weight (β) so that the final matrices followed an approximate scale-free topology (R2 ≥ 0.80). The dynamic tree-cut algorithm was used to detect network modules with a minimum module size set to 50 and cut tree height set to 0.99. These parameters were used to construct three separate networks. First, we built one large reference transcriptome network using all available mPFC samples from WT rats (*N*=35, genes=14,745, β=10). Nonspecific filtering required more than 2 counts per million (cpm) in at least 12 samples (1 batch) and retrained 14,552 transcripts. The resulting WT modules were assessed for enrichment for *Fmr1-^Δ^KH1* DGE signatures, FMRP targets, CNS cell type specificity, genetic risk loci for neurodevelopmental disorders and gene co-expression modules implicated in ASD cases. We then sought to determine whether any candidate WT modules displaying significant enrichment for *Fmr1-^Δ^KH1* DGE signatures and FMRP targets were also preserved in human cortex samples. To this end, we collected previously published healthy unaffected human cortical (BA 9/41) gene expression data [60] (RIN, 7.5 ± 0.6; Age, 33.25 ± 13.09; PMI, 23.58 ± 6.3; Sex, 16M/1F), and restricted our analysis to HGNC gene symbols that were commonly expressed in both rat and human data (genes = 6,926). Using this subset of genes, a second WT network was constructed (β=10) and a separate, third network was constructed for human cortical gene expression (β=15). Our module preservation analysis sought to determine whether any fundamental differences exist in the underlying gene co-regulatory patterns, as being preserved or disrupted, in WT rats as compared to humans, and *vice versa*. For these analyses, module preservation was assessed using a permutation-based preservation statistic, *Zsummary*, implemented within WGCNA with 500 random permutations of the data [101]. *Zsummary* takes into account the overlap in module membership as well as the density and connectivity patterns of genes within modules. A *Zsummary* score <2 indicates no evidence of preservation, *2<Zsummary*<10 implies weak preservation and *Zsummary* >10 suggests strong preservation.

### Functional annotation and protein interaction networks

Gene modules and DGE signatures with a FDR-corrected *P*-value <0.1 and an absolute log fold-change > 0.10 were subjected to functional annotation. First, the ToppFunn module [102] of ToppGene Suite software was used to assess enrichment of Gene Ontology (GO) terms specific to biological processes and molecular factors using a one-tailed hyper geometric distribution with family-wise FDR at 5%. Second, gene modules implicated in the neurobiology of FMR1 were used to build direct protein-protein interaction (PPI) networks, which can reveal key genes/transcription factors mediating the regulation of multiple target genes. PPIs were obtained from the STRING database [103] with a signature query of the reported module gene list. STRING implements a scoring scheme to report the confidence level for each direct PPI (low confidence: <0.4; medium: 0.4–0.7; high: >0.7). We used a combined STRING score of >0.7 and reported only the highest confidence interactions. We further used STRING to test whether the number of observed PPIs were significantly more than expected by chance using a nontrivial random background model (that is, null model). For visualization, the STRING network was imported into CytoScape [104].

### Module overlap and user-defined list enrichment analyses

DGE signatures and WT networks were annotated using previously defined gene sets. Cell type enrichment was performed by cross-referencing gene modules with previously defined lists of genes known to be preferentially expressed in different brain cell types [98]. Neurodevelopmental genetic risk loci were curated from human genetic studies of ASD [55–57], ID [58], SCZ [59] as well as dysregulated genes from mouse investigations [84, 105] and a list of well-known FMRP target genes [2]. Overrepresentation analysis of these gene sets within DGE signatures and WT transcriptome modules was analyzed using a one-sided Fishers exact test to assess the statistical significance. All *P*-values, from all gene sets and modules, were adjusted for multiple testing using Bonferroni procedure. We required an adjusted *P*-value < 0.05 to claim that a gene set is enriched within a user-defined list of genes. All user-defined lists can be found in **Supplementary Table 1**.

### Quantitative polymerase chain reaction (qPCR)

RNA was prepared is described above, from a new cohort of WT and *Fmr1-^Δ^KH1^−/y^* littermate rats (n=7/genotype). 1μg cDNA was synthesized from RNA samples, using the SuperScript II Reverse Transcriptase Kit (Invitrogen). The universal probe library (UPL) system (Roche) was used to perform quantitative polymerase chain reaction (qPCR). Two reference genes (*Rplp0* and *Gapdh*) were used for normalization. The relative expression levels were calculated using qBase software [106], now available from Biogazelle (Ghent, Belgium). Primers for each gene were designed using ProbeFinder Software (Roche). Primers location, primers sequences and UPL probe numbers are listed in **Supplementary Table 4**.

Acknowledgements
This work was supported by the Seaver Foundation (to JDB, HHN, SDR, CEMG, and MSB), Autism Speaks (to JDB), Autism Science Foundation Grant No. 17-001 (to MSB), and the grant F31 MH115656-01 (to CEMG). Carla E. M. Golden is a Seaver Graduate Fellow and Michael S. Breen is a Seaver Postdoctoral Fellow. We thank Eilam Doron, who contributed to this work.

